# Post-biting behavioral reprogramming underlies reproductive efficiency in *Aedes aegypti* mosquitoes

**DOI:** 10.1101/2025.06.25.661507

**Authors:** Linhan Dong, Elisabeth F. Bradford, Jord M. Barnett, Laura B. Duvall

## Abstract

The global spread and increasing populations of disease vector mosquitoes expose hundreds of millions of people to mosquito-borne illnesses each year. Female *Aedes aegypti* mosquitoes, global vectors of dengue, require protein from host blood to support egg development and undergo repeated cycles of blood-feeding and egg-laying. After biting, females temporarily alter their behavioral state and suppress host-seeking while using blood-derived nutrients to develop eggs. Host-seeking suppression ends once eggs are laid. While this period has generally been thought of as one of behavioral inactivity, we reveal that it instead reflects behavioral reprogramming, during which females transition from post-blood-meal inactivity into active searching for egg-laying sites. Females with mature eggs show a distinct behavioral state characterized by increased locomotor activity and a shift in circadian behavioral timing, leading to nocturnal humidity-seeking and egg-laying in an otherwise diurnal species. We show that the circadian clock gene *cycle* is critical for regulating this transition; its absence disrupts the timing of oviposition behaviors, leading to poor site selection and reduced predicted offspring survival. These findings suggest that during egg development, circadian clock-dependent behavioral reprogramming triggers nocturnal hyperactivity and oviposition site search, an essential process for mosquito reproduction and population viability.

## INTRODUCTION

To produce viable eggs, female *Aedes aegypti* mosquitoes must bite and obtain a blood meal from vertebrate hosts such as humans. This unique reproductive biology underlies their capacity to transmit disease-causing pathogens, including dengue, Zika, and chikungunya^1–4^. A single blood meal can double the female’s body weight, supplying the nutrients needed to develop dozens of eggs. This high reproductive output contributes to rapid mosquito population growth and global expansion^5–8^. Non-blood-fed females exhibit a strong host-seeking drive and are highly responsive to human-associated cues including carbon dioxide (CO_2_), human odor, and body heat to locate and bite hosts^9,10^. *Aedes aegypti* are primarily diurnal with peak biting activity and CO_2_ response persistence at dawn and dusk, and olfactory responses that vary throughout the day^11–15^. The circadian clock underlies the timing of nearly all biological rhythms and is governed by conserved clock genes that generate autoregulatory transcriptional-translational feedback loops^16–20^. Disruption of core clock genes, such as *cycle*, through mutation or RNAi, impairs circadian rhythmicity and alters mosquito behavioral patterns^21–24^. After blood-feeding to repletion, females undergo a dramatic behavioral shift: locomotor activity and responsiveness to host cues are suppressed for days during blood digestion and egg maturation^25–29^. During this period of host-seeking suppression, sensitivity to host-associated cues is reduced and, as eggs finally reach maturity, the gravid female transitions from inactivity to oviposition site selection^30–33^. Host-seeking suppression is reversible and typically ends after successful egg-laying, occurring 2–3 days post-blood-meal, although it can be prolonged if suitable oviposition sites are unavailable^34–37^. Previous studies in *Aedes* mosquitoes have reported variable peak oviposition times ranging from mid-afternoon to evening or night, but the role of the circadian clock in shaping these transitions remains untested^38–40^.

The search for an appropriate egg-laying site is crucial, as this location will serve as the habitat for the aquatic stages of a female’s offspring and ultimately determine her reproductive success^41,42^. Mosquito species exhibit niche-specific strategies when selecting egg-laying sites: some travel miles to reach natural freshwater reservoirs, while others specialize in locating artificial water containers near human dwellings or rain-filled objects in urban environments^35,43^. *Aedes aegypti* mosquitoes specialize in seeking freshwater sources around human habitats for egg-laying^44^. Gravid females rely on a combination of visual, humidity, and odorant cues to locate egg-laying sites^33,45–47^ and use their legs and proboscis to evaluate salinity and oviposition suitability^36^. Thus, following temporary post-blood-meal activity suppression, females must undergo a behavioral transition to initiate active searching and evaluation of oviposition sites. To investigate this shift, we used oviposition and activity-monitoring assays to characterize the behavioral states in anticipation of egg-laying. We found that *Aedes aegypti* females transition from quiescence to hyperactivity within three days of blood-feeding, coinciding with egg maturation, and initiate a distinct behavioral program that approximately doubles their daily activity. We further show that hyperactivity is closely linked to humidity-seeking, which is essential for effective oviposition site exploration and reproductive success. Notably, this hyperactive state diverges from the species’ typical diurnal activity, and instead reflects a nighttime pattern of egg-laying. The assignment of a nocturnal phase to egg-laying behaviors is lost in *cycle* (circadian clock) mutants. This unique gonadotrophic behavioral state demonstrates significant plasticity, with sensitivity to both internal and external environmental cues, and complete reversion upon successful oviposition.

## RESULTS

### Temporal reprogramming of activity in gravid females

After ingesting a blood meal, mated *Aedes aegypti* females suppress locomotor activity while they digest blood and develop eggs, which are ready to be deposited approximately three days later^25–27,29^ (**Figure 1A**). To characterize the behavioral state associated with egg-bearing females, we fed females a sheep blood meal and monitored locomotor activity over the following four days (**Figure 1B**). Consistent with previously described locomotor activity suppression^26,29^, females exhibited a pronounced reduction in overall activity during the first two days post-blood-meal, with a single peak occurring at dusk (**Figure 1B and 1C**). However, by the third day, coinciding with egg maturation, mosquitoes entered a hyperactive state with activity levels nearly double that of non-blood-fed controls, which persisted into the fourth day in the absence of an egg-laying site (**Figure 1C**). Despite this reactivation, the activity pattern remained unimodal with a prominent evening activity peak, differing from the bimodal rhythm observed in non-fed controls (**Figure 1B**). Notably, we also detected nocturnal hyperactivity in gravid females, which contrasts sharply with non-blood-fed females that show minimal activity at night (**Figure 1D**). The locomotor activity monitor does not provide an egg-laying substrate, confirmed by absence of eggs during the four-day recording period. To determine whether this hyperactive state is reversed upon oviposition, we allowed females to lay eggs in “ovi-vials” with a wet paper substrate (**Figure S1A**). After oviposition, females were transferred to locomotor activity monitors alongside age-matched, non-blood-fed controls. Activity levels and bimodal patterns were restored post-oviposition, confirming that the behavioral changes associated with egg-laying are reversible and state-dependent (**Figure 1E and F**, see also **Figure S1B**). Further, gravid females displayed no apparent phase shift compared to non-blood-fed controls under constant darkness (DD) conditions (**Figure S1C and D**), suggesting that this temporal reprogramming is likely not a direct result of a phase delay of internal circadian timing. Instead, we suspect that these behavioral changes are driven by downstream modulators of activity that alter the masking effect of darkness. Indeed, gravid females displayed similar amplitude of hyperactivity under both light-dark (LD) and DD conditions (**Figure S1E**). Overall, our data suggests that the gravid state temporarily overrides typical diurnal patterns and enables nighttime activity, even under dark conditions that normally suppress locomotion^24,48^.

**Figure 1.**
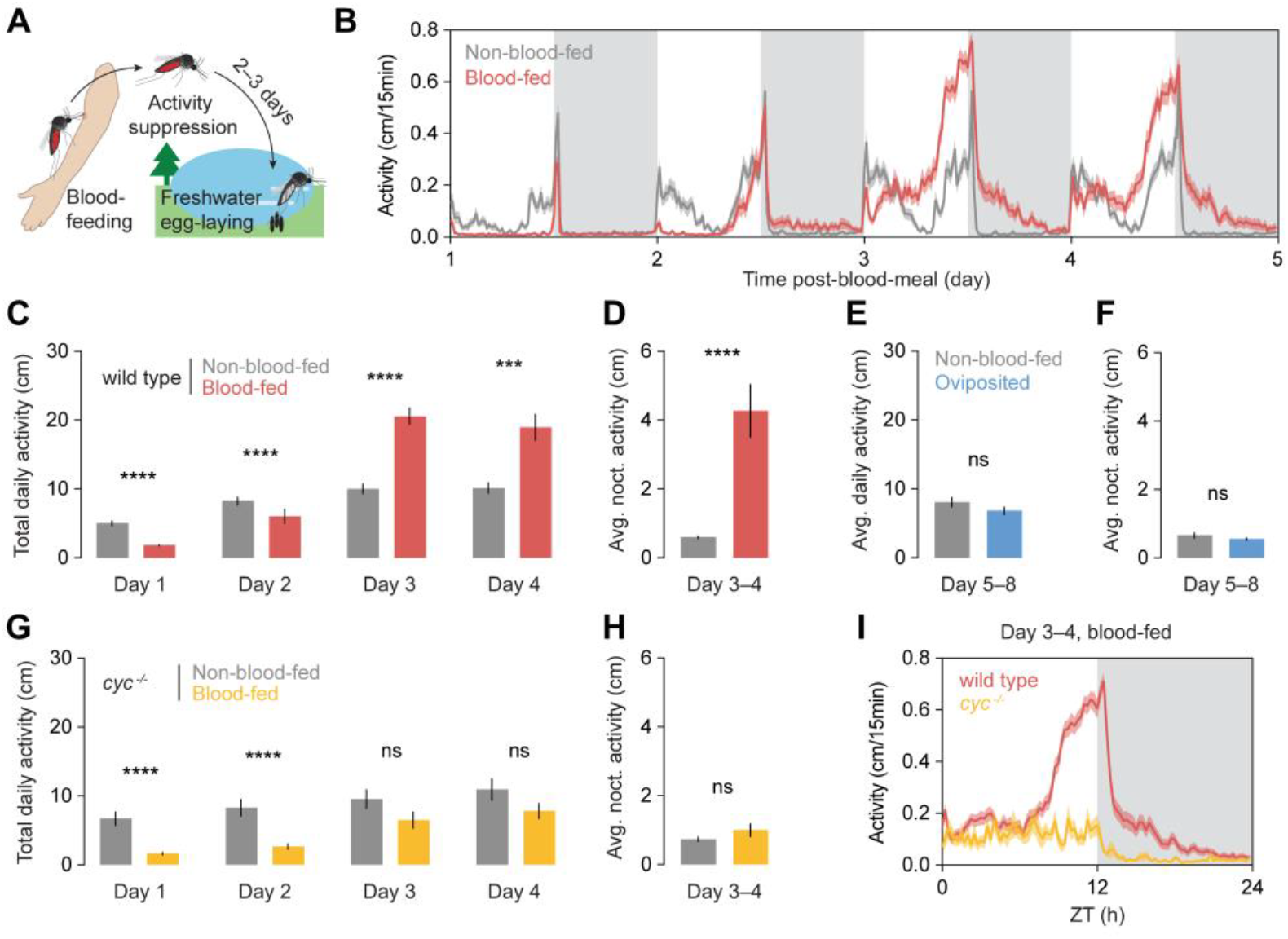
Alternative activity programming in gravid *Aedes aegypti* females is dependent on the clock gene *cycle*. (A) Schematic of the reproductive timeline of female *Aedes aegypti* mosquitoes. After feeding on a host, females begin blood meal digestion and egg development. Once eggs mature, females seek freshwater sites to lay eggs. (B) Locomotor activity profile of non-blood-fed (n = 58) and blood-fed (n = 56) *Aedes aegypti* mosquitoes. Locomotor activity (B–I) was recorded under 12 hours light: 12 hours dark LD cycles with 30 min dawn/dusk light transition periods. (C) Total daily activity of non-blood-fed (n = 58) and blood-fed (n = 56) *Aedes aegypti* mosquitoes. Mann-Whitney test. (D) Average nocturnal activity (ZT13–24) of non-blood-fed (n = 58) and blood-fed (n = 56) *Aedes aegypti* mosquitoes. Mann-Whitney test. (E) Average daily activity of non-blood-fed (n = 29) and oviposited (n = 18) *Aedes aegypti* mosquitoes. Mann-Whitney test. See also **Figure S1A and B.** (F) Average nocturnal activity (ZT13–24) of non-blood-fed (n =29) and oviposited (n = 18) *Aedes aegypti* mosquitoes. Mann-Whitney test. See also **Figure S1A and B.** (G) Total daily activity of non-blood-fed (n = 21) and blood-fed (n = 19) *cycle* mutants. Mann-Whitney test. (H) Average nocturnal activity (ZT13–24) of non-blood-fed (n = 21) and blood-fed (n = 19) *cycle* mutants. Mann-Whitney test. (I) Average daily activity profile of blood-fed wild type females (n = 56) and cycle mutants (n = 19), day 3 and 4 post-blood-meal. ZT: zeitgeber time, lights on at ZT0 and off at ZT12. White and grey shading in (B) and (I) represent light and dark phases, respectively. Lines and shaded lines in (B) and (I) show mean and SEM. Bars and error bars in (C–H) show mean and SEM. ***P < 0.001; ****P < 0.0001; P > 0.05: not significant (ns). Exact P values are provided in **Table S1**.

The circadian rhythmicity and synchrony of gravid females under DD conditions (**Figure S1D**) suggests that the clock temporally regulates this hyperactive state^49^. To determine whether a functional circadian clock is required for behavioral reprogramming in gravid females, we examined females with a loss-of-function mutation in the core clock gene *cycle*^23^. In these mutants, the disrupted CYC protein is unable to dimerize with its partner CLK, resulting in a disabled circadian clock and loss of bimodal activity under LD conditions^23^. In wild type non-blood-fed females housed under 12:12 LD conditions, evening activity persists briefly after lights-off, a pattern that is lost in *cycle* mutants (**Figure S2A**). Locomotor activity recording in DD conditions reveals decreased rhythmicity (**Figure S2B and C**), and *cycle* mutants exhibited drastically reduced locomotor activity in constant darkness, suggesting that masking effects of darkness are more dominant in the absence of a functional clock (**Figure S2B and D**). Following a blood meal, *cycle* mutants failed to exhibit the elevated activity levels and nocturnal hyperactivity observed in gravid wild type females (**Figure 1G and H**). These mutants also lacked anticipatory activity before dusk and did not show extended activity into the night (**Figure 1I**). These results demonstrate that the circadian clock, specifically the clock gene *cycle*, is essential for the emergence of the hyperactive, nocturnally-shifted behavioral state in gravid *Aedes aegypti* females.

### Nocturnal hyperactivity is temporally linked to humidity-seeking

While host-seeking remains suppressed during egg development, we reasoned that the hyperactive state in gravid females may reflect a shift in motivational state from seeking hosts to seeking suitable oviposition sites. *Aedes aegypti* are highly adapted to human environments, seeking out artificial water containers for egg-laying^44^. Additionally, they can exhibit “skip oviposition”, distributing eggs across multiple sites to maximize chances of offspring survival^50–53^. We hypothesize that hyperactivity facilitates oviposition site search specifically at night.

To test whether the observed activity shift is linked to sensory-driven oviposition behaviors, we examined humidity-seeking, as increased humidity level is an ecologically relevant cue that guides gravid females to freshwater egg-laying sites^46,47,54^. We developed a custom humidity preference assay in which mosquitoes were housed in a chamber with mesh windows above dry and water-containing Petri dishes while preventing direct contact with the water (**Figure 2A**). Humidity-seeking was quantified by comparing the number of dark pixel projections, indicating mosquito presence on the “humid” versus “dry” window (**Figure 2A**). Non-blood-fed females show consistently low humidity-seeking behavior. In contrast, gravid females show a rhythmic pattern with pronounced humidity-seeking in an approximately 5-hour window after dusk. This rhythm emerged three days post-blood-meal, coinciding with the onset of hyperactivity. In *cycle* mutants, although overall humidity preference increased after a blood meal, the rhythmic timing observed in wild type females was lost (**Figure 2B–D**). This suggests that proper timing of humidity-seeking requires a functional circadian clock, even in the presence of light-dark cycles. These results show strong temporal coupling between nocturnal activity and humidity-seeking in gravid females, indicating that the circadian system plays a critical role in coordinating the timing of oviposition-related behaviors to ensure that egg-laying occurs at an optimal time of day.

**Figure 2.**
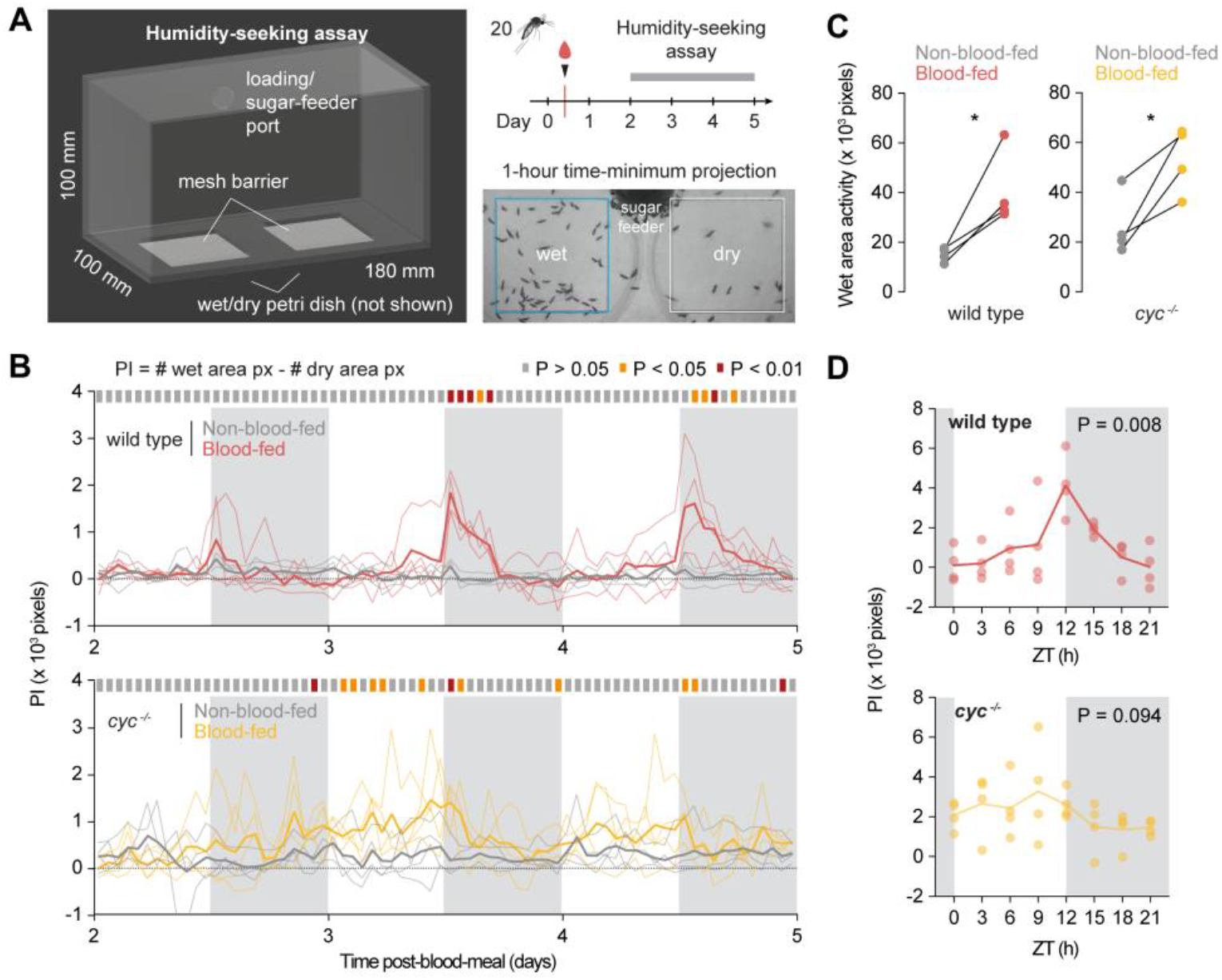
Rhythmic, nocturnal humidity-seeking is regulated by the clock gene *cycle*. (A) Schematic of the humidity-seeking assay and experimental timeline. (B) Humidity-seeking preference index (PI) of wild type and *cycle* mutant females (blood-fed and non-blood-fed). For (B–D), n = 4 biological replicates with 20 females per replicate. Unpaired t-test. (C) Cumulative humidity-seeking activity of wild type and *cycle* mutant females (blood-fed and non-blood-fed, day 3–4 post-blood-meal). Paired t-test. (D) Average humidity-seeking profile of blood-fed wild type and *cycle* mutant females, day 3–4 post-blood-meal. BioDare2 eJTK rhythmicity test. White and grey shading in (B) and (D) represent light and dark phases, respectively. Thick lines in (B) and (D) show mean, and thin lines in (B) and dots in (C) and (D) show independent replicates. *P < 0.05. Exact P values are provided in **Table S1**.

### Nocturnal egg-laying in a diurnal mosquito

In non-blood-fed *Aedes aegypti* mosquitoes, daily behavioral timing follows a diurnal, bimodal pattern with peaks at dawn and dusk in laboratory and field studies^11,13–15,55^. Our data suggest that gravid females exhibit an alternative circadian program characterized by nocturnal hyperactivity and humidity-seeking that may reflect the preferred time of day for egg-laying. *Aedes aegypti* mosquitoes require contact with a wet surface to lay eggs (**Figure 3A**)^43,45^. While previous studies have explored oviposition rhythms by manually replacing egg-laying substrates every few hours, this approach is labor-intensive and susceptible to unwanted experimenter influence^39,40,53,56,57^. Across insect species, monitoring the internal timing of oviposition remains a technical challenge and efforts have focused on developing automated systems to generate oviposition rhythms with minimal experimenter bias^58^. Here, we introduce “EggTimer,” a behavioral monitoring platform that enables continuous automated tracking of egg-laying (**Figure 3B**). In this setup, mosquitoes are individually housed above a wet paper substrate connected to a water reservoir and illuminated with infrared (IR) light, to allow real-time visualization of egg deposition and locomotor activity. *Aedes aegypti* eggs are initially white due to their transparent endochorion, but melanize and darken within three hours after being laid^41,59–61^. We developed a custom image-processing algorithm to detect pixel-level changes in darkness, enabling accurate determination of oviposition timing (**Figure 3C**). Using this system, we observed that, despite their diurnal locomotor activity prior to blood-feeding, *Aedes aegypti* females displayed a unimodal oviposition rhythm when given continuous access to egg-laying substrate, with oviposition occurring most frequently immediately after dusk (**Figure 3D and E**). In *cycle* mutants, rhythmicity and nocturnal preference is disrupted (**Figure 3D and E**), and eggs are laid throughout the day, demonstrating that circadian control is essential for proper oviposition timing.

**Figure 3.**
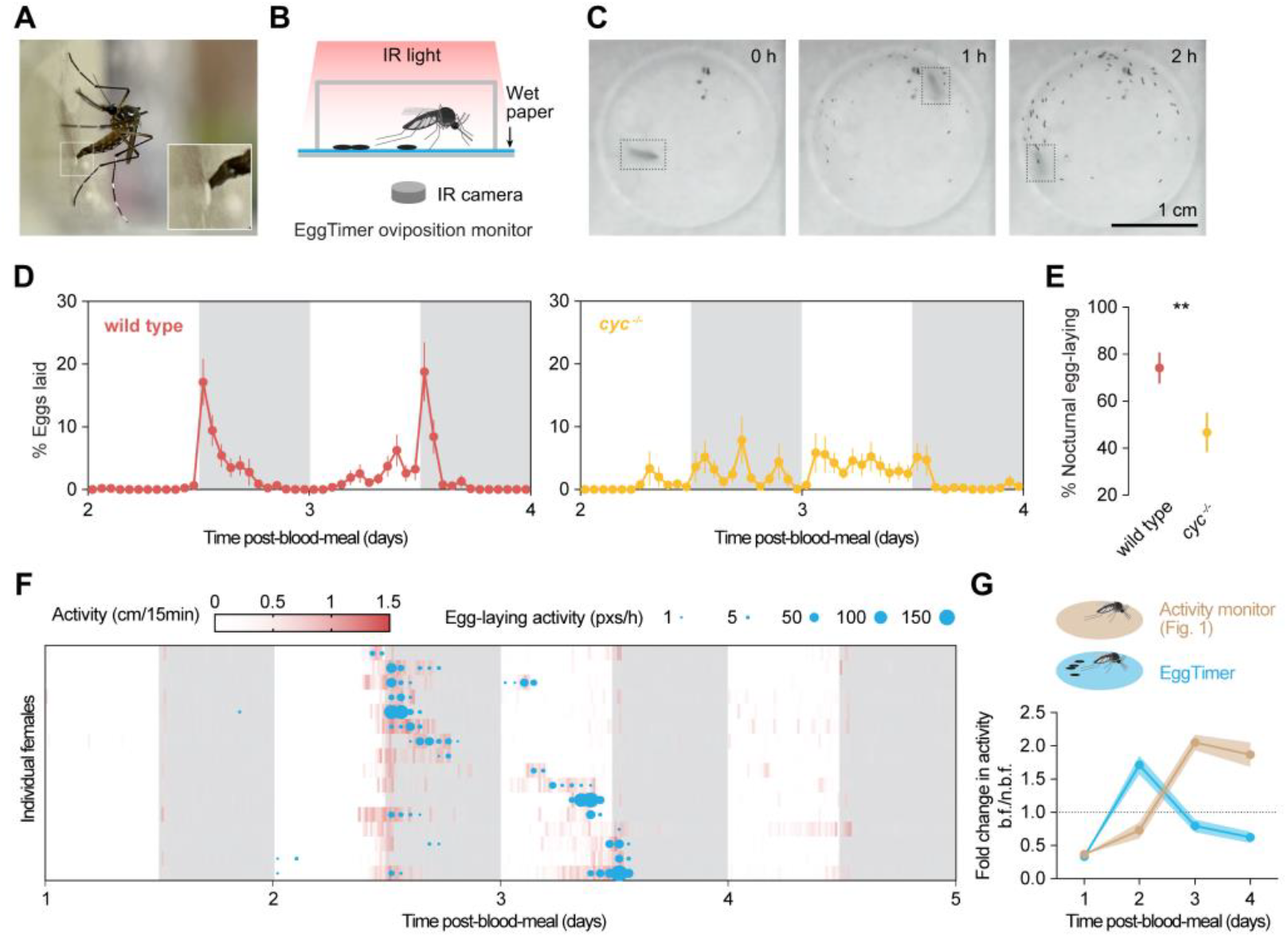
Hyperactivity is contingent upon egg-retention. (A) A gravid female *Aedes aegypti* mosquito laying eggs on a wet paper substrate. Newly laid eggs are white and darken over time. (B) Schematic of the EggTimer oviposition monitor. (C) Representative images of an egg-laying female acquired with the EggTimer oviposition monitor. Boxes indicate the shadow of the female mosquito. (D) Egg-laying rhythm in wild type (n = 41) and *cycle* mutant (n = 30) females. (E) Percentage of eggs laid in scotophase (ZT12–24). Mann-Whitney test. (F) Representative egg-laying and locomotor activity of wild type females in the EggTimer assay. Individuals shown from one cohort. (G) Fold change in daily activity comparing blood-fed and non-blood-fed wild type females in the locomotor activity monitor (n = 56) and the EggTimer (n = 38). White and grey shading in (D) and (F) represent light and dark phases, respectively. Dots and error bars/shading in (D), (E), and (G) show mean and SEM. **P < 0.01. Exact P values are provided in **Table S1**.

In this assay, gravid females typically laid eggs between the second and third day post-blood-meal, suggesting that when housed directly on an egg-laying substrate, hyperactivity onset may occur earlier than when no suitable oviposition site is available. To test this, we conducted extended EggTimer recordings on blood-fed females using an experimental timeline that matched our locomotor activity recordings (**Figure 1B**). We tracked locomotor activity using the mosquito’s shadow on the wet paper substrate to infer the female’s activity level and temporal relationship to oviposition (**Figure 3C**; see also **Figure S3A**). By the second day post-blood-meal, females showed a reactivation of activity that was tightly aligned with the onset of egg-laying (**Figure 3F**). Notably, both overall and nocturnal hyperactivity peaked on the day of oviposition but returned to baseline the following day (**Figure 3F**; see also **Figure S3B**). To assess whether the presence of an oviposition substrate influenced the magnitude or timing of hyperactivity, we compared locomotor activity over four days in two conditions: females housed with an oviposition substrate (EggTimer assay) and females without access to oviposition substrate (standard locomotor activity assay, **Figure 1**). Females without substrate showed sustained hyperactivity beginning on day 3 post-blood-meal, while those with substrate exhibited earlier hyperactivity on day 2 that resolved by day 3 (**Figure 3G**). This suggests that access to a substrate reduces the need for prolonged movement (see also **Figure S3C**). Nonetheless, a nocturnal shift in activity persists and remains tightly coupled to nighttime egg-laying. These findings indicate that the hyperactive state is internally driven but modulated by environmental context.

### Hyperactivity supports long-range oviposition site-seeking

Gravid females must navigate and evaluate a complex and variable sensory landscape when selecting oviposition sites in human environments^62^. Suboptimal or hazardous sites, such as those with high salinity, can severely compromise reproductive success. If suitable sites are not found within a few days of egg maturation, females risk laying eggs in lethal conditions or may fail to lay them altogether^34–37^. Our data suggest that gravid females enter a hyperactive behavioral state with altered activity timing to maximize the likelihood of successful oviposition – the final and critical step of the reproductive cycle. Without a functional clock, *cycle* mutants do not exhibit hyperactivity and nocturnal humidity-seeking, and we hypothesize that this disrupts their ability to search for and locate suitable oviposition sites. To model a scenario requiring extended search behavior, we designed an assay consisting of five sequentially connected cages: the first three cages next to the starting cage each contained an ovi-cup filled with 0.24 M NaCl (lethal to larval growth^36^), while the final cage contained an ovi-cup with freshwater (**Figure 4A**). We released a group of 20 gravid females in the starting cage at ZT11.5 (30 minutes before dusk) of the third day post-blood-meal and allowed them to explore and lay eggs for the next five hours, coinciding with peak humidity-seeking and egg-laying observed in our earlier assays. To ensure that differences in egg-laying were due to exploratory and not physiological deficits, we confirmed that *cycle* mutants lay eggs as efficiently as wild type females when housed directly on a freshwater oviposition substrate and tested each genotype separately to ensure that community-based cues did not influence site selection^63^ (**Figure S4A and B**).

**Figure 4.**
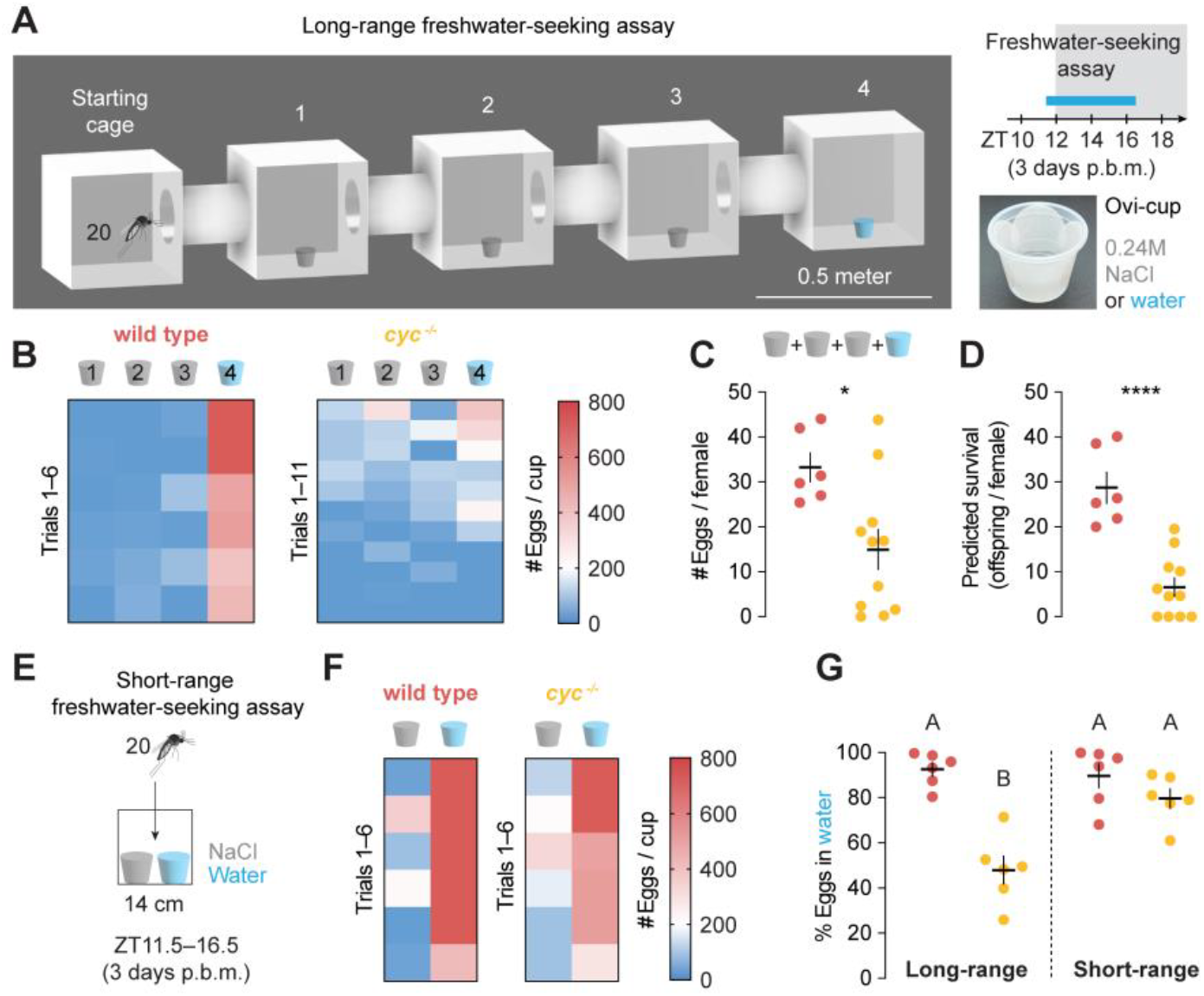
Hyperactivity underlies reproductive efficiency. (A) Schematic of the long-range freshwater-seeking behavioral assay and ovi-cups used for egg collection. Animals three days post-blood-meal (p.b.m.) were released in groups of 20 at ZT11.5 and allowed five hours in the behavioral assay. (B) Heatmap of egg-laying results by wild type (n = 6) and *cycle* mutant (n = 11) *Aedes aegypti* mosquitoes in the long-range freshwater-seeking assay. (C) Eggs laid per female across all cups by wild type (n = 6) and *cycle* mutant (n = 11) females in cage 1–4 of the long-range freshwater-seeking assay. Unpaired t-test. (D) Number of surviving offspring per female predicted with oviposition-site lethality by wild type (n = 5) and *cycle* mutants (n = 11). Unpaired t-test. (E) Schematic of the short-range freshwater-seeking behavioral assay. Animals three days p.b.m. in groups of 20 were released at ZT11.5 and allowed five hours in the behavioral assay. (F) Heatmap of egg-laying results by wild type (n = 6) and *cycle* mutants (n = 6) in the short-range freshwater-seeking assay. (G) Freshwater preference of wild type and *cycle* mutants in the long-vs short-range freshwater seeking assay (n = 6 for all conditions and genotypes). Trials with less than eight eggs per female were excluded to ensure sufficient participation. Ordinary one-way ANOVA with Tukey’s multiple comparisons test. Data labeled with different letters are significantly different from each other (P < 0.001). Line and error bars in (C), (D), and (G) show mean and SEM and dots show individual trials. *P < 0.05; ****P < 0.0001. Exact P values are provided in **Table S1**.

In the long-range assay, wild type females laid eggs almost exclusively in the farthest freshwater cup, showing a strong avoidance of the lethal substrates (**Figure 4B**). *Cycle* mutants, however, deposited more eggs in the lethal cups and showed reduced egg deposition in the freshwater option (**Figure 4B**). Despite similar egg-laying efficiency when placed directly on an oviposition substrate, *cycle* mutants laid significantly fewer eggs overall in the long-range freshwater-seeking assay, likely due to a lack of hyperactivity required to access oviposition sites (**Figure 4C**). This suggests that reduced egg output is not due to general impairment in egg production but may be attributable to defects in egg-laying timing (**Figure 3D**) and site-searching in the long-range assay. The combination of limited exploratory behavior and inappropriate oviposition in lethal sites results in significantly lower reproductive success in *cycle* mutants, measured by predicted surviving offspring^36^ (**Figure 4D**). To investigate whether *cycle* mutants show impairments in detecting salinity or making oviposition choices on a small spatial scale, we established a short-range freshwater-seeking assay (**Figure 4E**). In this short-range setup (40 times smaller than the long-range assay), *cycle* mutants consistently laid more eggs in the freshwater cup (**Figure 4F**). We then calculated freshwater preference in all trials with greater than 8 eggs per female. The defect in freshwater preference of *cycle* mutants in the long-range assay was recovered in a short-range setup, indicating that they can recognize suitable oviposition sites at close range (**Figure 4G**). We hypothesize that the impaired long-range oviposition behavior in *cycle* mutants results from a combination of increased internal pressure to lay eggs and reduced exploratory behavior, due to the absence of nocturnal hyperactivity, ultimately compromising their ability to locate optimal egg-laying sites. Indeed, when no suitable oviposition substrate is available, wild type females will eventually deposit eggs in a saltwater cup, although with reduced efficiency and avoidance of the paper substrate (**Figure S4C and D**).

## DISCUSSION

While circadian regulation is typically defined by robust, self-sustained rhythms under constant conditions, examples of plasticity in daily behavioral timing that result in temporal niche switches are found across the tree of life^64^. These shifts can be triggered by environmental and physiological factors including energy balance^65,66^ and seasonal changes^67,68^. In this study, we characterize a striking example of temporal niche flexibility in the diurnal mosquito *Aedes aegypti*, which exhibits a transient, dusk-to-nighttime preference for egg-laying. This oviposition timing mirrors patterns observed in nocturnal mosquito species such as *Anopheles gambiae*^69^, despite *Aedes aegypti* being otherwise day-active. The convergence of oviposition timing across both diurnal and nocturnal species points to the existence of evolutionarily conserved circadian mechanisms that coordinate reproductive timing across diverse mosquito lineages and suggests that nighttime egg-laying may confer general adaptive advantages. Newly-laid eggs lack a melanized endochorion, making them more susceptible to UV radiation and desiccation^59^. Laying eggs at sunset may offer optimal environmental conditions including lower temperature, higher humidity, and diminished UV level that promote egg viability. Our findings show that *Aedes aegypti* females, although typically diurnal, are capable of reprogramming their activity to adopt a nocturnal egg-laying schedule. This behavioral reprogramming requires the molecular circadian clock. In *cycle* mutants, the rhythmicity and timing of both humidity-seeking and oviposition are disrupted, even under entraining light-dark conditions, indicating that an intact molecular clock is essential for coordinating these behaviors. Supporting this, gravid wild type females maintain circadian rhythmicity under constant dark conditions, suggesting that the behavioral changes arise from clock-driven output pathways. Intriguingly, “restless” egg-laying nights in gravid females draw parallels with seasonal behavioral changes in migratory birds known as “zugunruhe” or nocturnal restlessness observed in otherwise diurnal birds^70^. This circannual migratory drive is conserved in nonmigratory birds^71^, suggesting a potential utility of this alternative program in migratory flexibility in variable environments. However, unlike seasonal niche switches, the shift to nocturnal activity in mosquitoes is transient, tightly coupled to reproductive physiology, and reversed immediately following egg deposition. The features we observe in female mosquitos define a unique form of temporal niche plasticity that is both specific to egg-bearing females and under strong circadian control. Such plasticity highlights the adaptability of behavioral timing in the service of reproductive success.

Accompanying the shift in behavioral timing, gravid females enter a hyperactive state that overrides post-blood-meal activity suppression. Given that this heightened activity is sustained during egg retention and rapidly ceases following oviposition, we propose that it represents a short-term, energetically costly strategy that enhances the likelihood of locating suitable egg-laying sites, which ultimately maximizes reproductive success. Mosquito blood-feeding occurs across a range of environments varying in urbanization, vegetation, and microclimate^35,43^. While some mosquito species undertake long-range migrations in search of oviposition sites, *Aedes aegypti* is more commonly associated with human-modified habitats, actively seeking out small artificial containers such as tires, pots, and gutters^32,44,54,72^. In such unpredictable landscapes, hyperactivity may enable gravid females to detect and integrate a wide array of sensory cues—physical, chemical, and visual that vary in signal strength and range. Indeed, previous descriptions of mosquito oviposition behavior refer to a phase of seemingly “random activity,” thought to aid in the discovery of suitable sites^45,54^. Our findings support this notion, highlighting hyperactivity as an adaptive behavioral state that facilitates site evaluation under uncertain environmental conditions. The rapid cessation of hyperactivity post-oviposition suggests that signals from the gravid ovary may sustain this elevated activity state, and earlier studies have shown that transplanting a gravid ovary into a non-blood-fed female can elicit pre-oviposition behaviors, implicating ovarian signals in modulating behavior^73^. Future work aimed at identifying the molecular nature of these signals could uncover novel targets to reduce reproductive efficiency.

The unique reproductive biology of the female mosquito provides a compelling model to investigate the innate regulation of behavior. Our study describes distinct internal states that define host-seeking, blood digestion, and egg-laying phases that are accompanied by shifts in sensory tuning and circadian timing. Together, this sequence of transitions reveals an extraordinary degree of behavioral plasticity. Identifying the mechanisms that control behavior across these key reproductive stages will advance our understanding of mosquito biology and opens new approaches for vector control. These results demonstrate that the circadian clock is essential for the behavioral reprogramming that supports efficient egg-laying.

Targeting this tightly regulated gonadotrophic cycle by disrupting the circadian system could provide novel approaches to prevent mosquito reproduction and to mitigate disease transmission.

## ACKNOWLEDGEMENTS

We thank Thomas Gabel for assistance with animal husbandry and Vinaya Shetty and Zach Adelman for providing the *cycle* mutant line. We thank Nils Reinhard, Ben Matthews, Paul Taghert, and members of the Duvall lab for useful discussions and feedback on the manuscript. This work was supported by the following grants: NIGMS (R35 GM137888) (LBD), Beckman Young Investigator Award (LBD), Pew Scholar in Biomedical Sciences Award (LBD), Klingenstein-Simons Fellowship Award in Neuroscience (LBD).

## AUTHOR CONTRIBUTIONS

LD carried out or supervised all experiments and data analysis with contributions from coauthors. EFB designed and carried out egg-laying assays in **Figure 4** and **Figure S4**. JMB assisted with piloting and data analysis of the EggTimer assay in **Figure 3**. LD, EFB, and LBD together conceived the study, designed the figures, and wrote the paper with input from all authors.

## SUPPLEMENTAL INFORMATION

**Table S1**. All raw data and statistical test details used for main and supplemental figures.

## METHOD DETAILS

### Mosquito rearing

*Aedes aegypti* Liverpool mosquitoes and *cycle*^*23*^ mutant strains were maintained and reared at 28°C, 70–80% relative humidity with a 12 h light: 12 h dark schedule. Eggs were hatched using 1 L of TetraMin fish food suspension in deionized deoxygenated water (one tablet/L). Larvae were fed with TetraMin fish food tablets each day as needed. Adult female mosquitoes used for experiments were 3–21 days old and co-housed with males, with *ad libitum* access to 10% sucrose solution until further experimental procedures. For passaging, adult female mosquitoes were blood-fed on defibrinated sheep blood (Quad5) with 1 mM ATP at 37°C using Hemotek artificial membrane feeders.

### Blood-feeding for behavioral assays

All animals tested for post-blood-meal behavior were entrained for at least three days under a 12 h light: 12 h dark schedule. Access to sucrose solution was removed and a cotton wick soaked with DI water was provided approximately 24 hours before blood-feeding. Animals were allowed to blood-feed on defibrinated sheep blood (Quad5) with 1 mM ATP at 37°C using Hemotek artificial membrane feeders. The feeder was rubbed on experimenter’s arms to provide human odor. The experimenter’s breath was used to improve feeding rate. After 30 minutes of access to a feeder, cages were transferred to 4°C to cold anesthetize animals. Females were scored by eye for complete abdominal distension indicating that they fed to repletion, non-fed animals were removed, and fully blood-fed animals were returned to 28°C, 70–80% relative humidity. All feeding was conducted at ZT8–9.

### Locomotor activity assay

Locomotor activity was assessed using a modified version of the MozzieBox locomotor activity assay^11^. This modified design is constructed with four 23 cm miniature t-slotted framing (McMaster-Carr) and custom cut acrylic sheets (3 mm thickness, black or clear) that has overall dimensions of 29 × 18 × 31 cm to fit inside a compact incubator (Heratherm, Thermo Fisher) to allow for temperature control. All recordings were performed with incubator temperature set at 28°C. Infrared LED lights (850nm, Waveform Lighting) were installed at the bottom panel, and four 4 × 4 cm cooling fans (Noctua NF-A4×10 FLX) were installed to improve heat dissipation and air circulation. RaspberryPi Camera Module 3 NoIR Wide angle cameras covered with IR longpass filters (Edmund Optics 43-948) were mounted on top of the acrylic frames to take images every minute. White LED strips were installed and controlled together with the camera via custom Python script. All platforms and components were assembled with corner brackets and surface brackets for miniature t-slotted framing (McMaster-Carr). Python scripts can be found on Github (https://github.com/DongLinhan/EggTimer2025).

Each well of a 12-well tissue culture plate was supplied with 1.5 mL of 10% sucrose in 1% agarose, with 1% (v/v) 15% p-hydroxy-benzoic acid methyl ester in 95% ethanol (antifungal). Female mosquitoes (non-blood-fed, blood-fed, oviposited) were individually loaded into the wells under cold anesthesia. All recordings of blood-fed or oviposited females were performed with age-matched controls from the same rearing cage. Loading of blood-fed animals was conducted immediately after blood-feeding at ZT8. For post-oviposition activity recording, females were loaded into individual ovi-vials on day two post-blood-meal, constructed using *Drosophila* vials (VWR 75813-164), folded filter paper (55 mm, Whatman), sitting in 5 mL of DI water. Animals were only loaded for post-oviposition activity recording if a full clutch of over 50 eggs was laid in the ovi-vial by day four post-blood-meal. Up to four plates were loaded into the monitor before ZT10, and recording started the day after at ZT0. For all LD recording, four LD cycles with 30 min dawn and dusk periods were applied and IR images were taken every minute. For recordings in **Figure S1C**, one LD cycle was applied before three days of DD conditions. For recordings in **Figure S2**, three LD cycles were first applied, before switching to DD for seven days. Downstream analysis of images uses the same pipeline as in Dong et al. (2024)^11^. Briefly, images were converted into a 30 frames-per-second video using a custom Matlab code, and the video was further analyzed using Ethovision (version 17) to extract positional features for movement analysis. Analysis of circadian rhythmicity and phase synchrony used the same pipeline and custom coding as in Dong et al. (2024)^11^. Briefly, raw distances were transformed into activity scores that represents activity in one-minute bins. Calculation of rhythmic power were performed using the phase package (1.2.9) in R (version 4.2.3)^74^. Animal is considered rhythmic if rhythmic power is greater than 50. For phase quantifications of DD activity, a Savitzky-Golay filter was applied to animal activity scores in the third and fourth day of recording and the phase was averaged between the two days for further comparison. Phase synchrony was determined using the Rayleigh R test.

### Humidity-seeking assay

Humidity-seeking activity was assessed using the MozzieMonitor platform^75^ with custom-built humidity-seeking arenas (**Figure 2A**). The humidity-seeking arena is a 10 × 10 × 18 cm acrylic box with two 5 × 5 cm open mesh windows constructed with 3 mm clear acrylic sheets. Facing the mesh windows, a round Petri dish (100 mm x 15 mm, Celltreat) filled with DI water and an empty Petri dish were placed directly under the arena. Sucrose solution was provided to the arenas via a cotton wick partially soaked in 10% sucrose. For each experiment, 20 blood-fed females and 20 non-blood-fed control females were aspirated into two separate behavioral arenas one day after blood-feeding. Animals were allowed to acclimate in the behavioral arena and image acquisition began the following day at ZT0 for three days under LD conditions with 30 min dawn and dusk periods. IR images were taken every minute, and a time-maximum projection was created using custom Python code as background. Every 60 images were converted into a time-minimum projection, and pixels were counted if below darkness threshold (less than 100). Pixel counts were subtracted from the background image and grouped by user-defined ROI using custom Python coding. The analysis code used can be found on Github (https://github.com/DongLinhan/EggTimer2025). Rhythmicity test of humidity seeking activity was carried out using BioDare2^76^ (https://biodare2.ed.ac.uk).

### EggTimer oviposition monitor

The EggTimer assay was modified from the MozzieMonitor platform^75^ with adaptations on platform heights and orientation. A RaspberryPi Camera Module 3 NoIR Wide camera was mounted on the bottom platform covered with an IR longpass filter (Edmund Optics 43-948). Miniature t-slotted framing (30 cm, McMaster-Carr) was installed 11 cm above the camera to support a removable platform made with 3 mm acrylic sheet. An IR lighting platform was installed 9 cm above this removable platform, and two 14 × 14 cm USB fans (Newegg 9SIB7W5HBZ4935) were added to the IR lighting platform to provide ventilation and prevent overheating. An additional acrylic sheet covered in Kimwipe was installed 1.5 cm below the IR lights to provide diffusion of the IR light. Two open plastic containers (6 × 8 × 13.5 cm) filled with DI water were placed on the camera platform to provide a water source for the wet paper substrate. Up to two 12-well tissue culture plates can be imaged on the imaging platform. The imaging platform is covered in two layers of wet Kimwipes, leaving overhangs on each side of the platform to be soaked in the water source. For oviposition recordings, blood-fed females were loaded into individual wells of a 12-well tissue culture plate one day after blood-feeding under cold anesthesia. Animals were allowed to acclimate in the behavioral arena and image acquisition began the following day at ZT0 for three days under LD conditions with 30-min dawn and dusk periods. For long-term oviposition/locomotion recordings, loading of blood-fed animals was conducted immediately after blood-feeding at ZT8 to match the timeline of the locomotor activity recordings. Images were taken every minute and female movement analysis follows the same pipeline as the locomotor activity assay. For egg-laying activity, we established a pipeline to detect pixel value change overtime that is characteristic of a darkening egg. While white, un-melanized eggs is visible and induce pixel value change under IR imaging, we used the pixel darkening profile to verify this initial change to reduce noise signals from animal movement. Pixel values were extracted from images acquired during recordings and time series of pixel values were processed with a Savitzky-Golay filter (window size = 15, polynomial order = 3). For each individual time series of pixel values, two rolling windows (A: 30 minutes, B: 240 minutes) were used to detect a significant and smooth drop in pixel value that persists until the end of the recording. The end of window A was then recorded as the egg-laying onset if window A and B matched the numerical criteria established in our algorithm. Detailed analysis code, along with data processing procedures and corresponding threshold values are deposited on Github (https://github.com/DongLinhan/EggTimer2025).

### On-substrate assay

Egg-laying behavior was assessed by housing females three days post-blood-feeding directly in ovi-cups, eliminating the need for long-range oviposition site-seeking. A 4-oz cup with cap (VWR 8900-662) was filled with 50 ml DI water or 0.24 M NaCl and four pieces of filter paper (55mm, Whatman). A 35 × 10 mm plastic Petri dish (Greiner) was fastened to the inner side of the ovi-cup cap with screws and was used to house females until the start of the experiment. Before each experiment, 10 females were cold anesthetized and placed in the Petri dish inside the ovi-cup, and returned to 28°C, 70–80% relative humidity to acclimate for 90 minutes. At ZT11.5, the Petri dish cap was lowered by 1 cm, releasing females into the arena. At ZT13, ovi-cups were moved to a cold room to anesthetize and remove females, concluding the experiment. All handling was done with gloves to avoid human odor contamination. For saltwater trials, eggs were manually counted. For freshwater trials, images of the eggs were taken using a webcam (Logitech) and dark pixels were converted into egg counts. Analysis code can be found on Github (https://github.com/DongLinhan/EggTimer2025).

### Long-range freshwater seeking assay

Long-range freshwater-seeking behavior was assessed in an arena consisting of five 30 × 30 × 30 cm cages (BugDorm DP1000) connected via custom 20 cm mesh tunnels with openings 15 cm in diameter (BugDorm DP1000-N) (**Figure 4A**). All assays were performed in an environmentally-controlled room at 28°C, 70–80% relative humidity. Animals were tested three days post-blood-feeding and 20 females were moved to an 8-oz plastic cup with a cap (VWR 8900-664) under cold anesthesia.

Females were placed in the starting cage to acclimate for 90 minutes before experimentation. Ovi-cups were made with 4-oz cups (VWR 8900-662) with four pieces of filter paper (55mm, Whatman) lining the sides and filled with 50 ml 0.24 M NaCl solution, which is lethal to mosquito larvae^36^, or DI water. Salt ovi-cups were placed in cages 1–3 (closest to starting cage), and the freshwater ovi-cup was placed in cage 4 (farthest from starting cage). Handling of arena components and mosquitoes was done with gloves to avoid contamination with human odor. At ZT11.5, females housed in the 8-oz cup were released into the arena and allowed five hours in the behavioral setup. Ovi-cups were removed at ZT16.5 and eggs laid in ovi-cups were allowed to melanize overnight and manually counted the following day. To report preference between salt and freshwater, we excluded trials with fewer than 150 eggs total collected to prevent skewed preference due to low participation. Predicted survival was calculated using published data^36^, where freshwater substrate led to 92.22% survival and 0.24 M NaCl used in this study is above the threshold concentration that leads to immediate larval lethality.

### Short-range freshwater seeking assay

Short-range freshwater-seeking behavior was assessed three days post-blood-feeding, and 20 females were aspirated into a 14 × 14 × 14 cm small mesh cage (BugDorm 4M1515) and acclimated for 90 minutes. Two 4-oz ovi-cups (VWR 8900-662) filled with either 50 ml DI water or 50 ml 0.24 M NaCl with four pieces of filter paper (55mm, Whatman) lining the sides were placed in the cage at ZT11.5. Experiments were concluded at ZT16.5 by removing the ovi-cups and eggs laid in ovi-cups were allowed to melanize overnight and manually counted the following day.

## Statistics

All statistical analyses were performed with GraphPad Prism or custom Python code. Statistical methods and sample sizes are included in figure captions. Nonparametric tests were used for data that does not follow normal distribution. Source data and statistical test details, including exact n and P values, can be found in **Table S1**.

## SUPPLEMENTAL FIGURES

**Figure S1.**
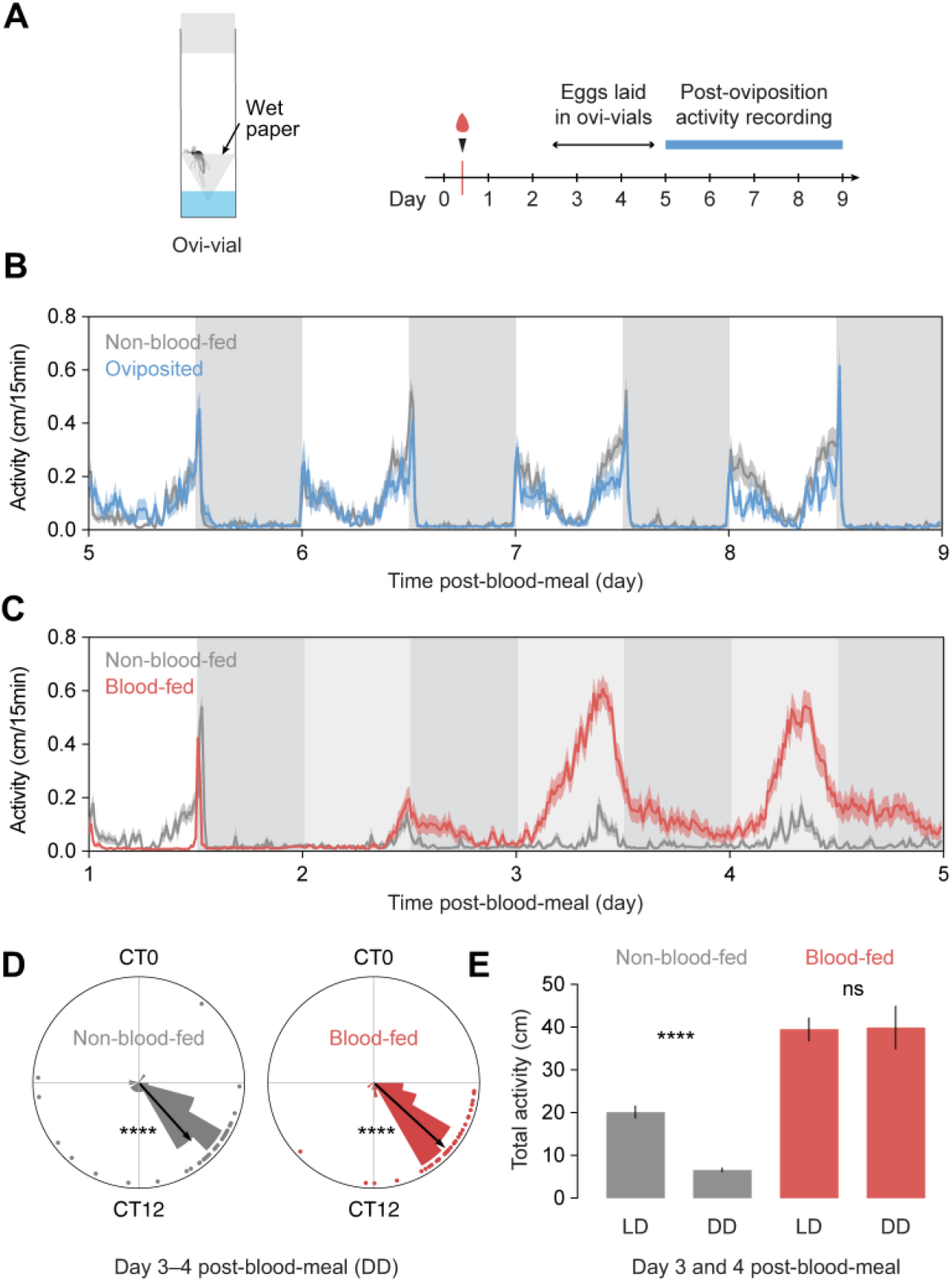
Hyperactive behavioral programming in *Aedes aegypti* is reversible and under circadian control. (A) Schematic of ovi-vial and experimental timeline of post-oviposition activity recordings. (B) Locomotor activity profile of non-blood-fed (n = 29) and oviposited (n = 18) *Aedes aegypti* mosquitoes under LD entrainment. (C) Locomotor activity profile of non-blood-fed (n = 34) and blood-fed (n = 40) *Aedes aegypti* mosquitoes under one day of LD entrainment, followed by three days of DD entrainment. (D) Circular plots of the phase of activities in day 3–4 post-blood-feeding under DD (C). The vectors (arrows) pointing from the center towards the perimeter show the average peak activity time and vector length corresponds to the extent of phase coherence. Asterisks correspond to the P-value of Rayleigh R test for phase coherence. (E) Total activity days 3–4 post-blood-feeding under LD (Figure 1B) and DD conditions (C). Unpaired t-test. White and grey shading in (B) and (C) represent light and dark phases, respectively. Light grey shading in (C) represents respective light phase from prior LD entrainment. Lines and shaded lines in (B) and (C) show mean and SEM. Dots in (D) shows individual animals. Bars and error bars in (E) show mean and SEM. ****P < 0.0001; P > 0.05: not significant (ns). Exact P values are provided in **Table S1**.

**Figure S2.**
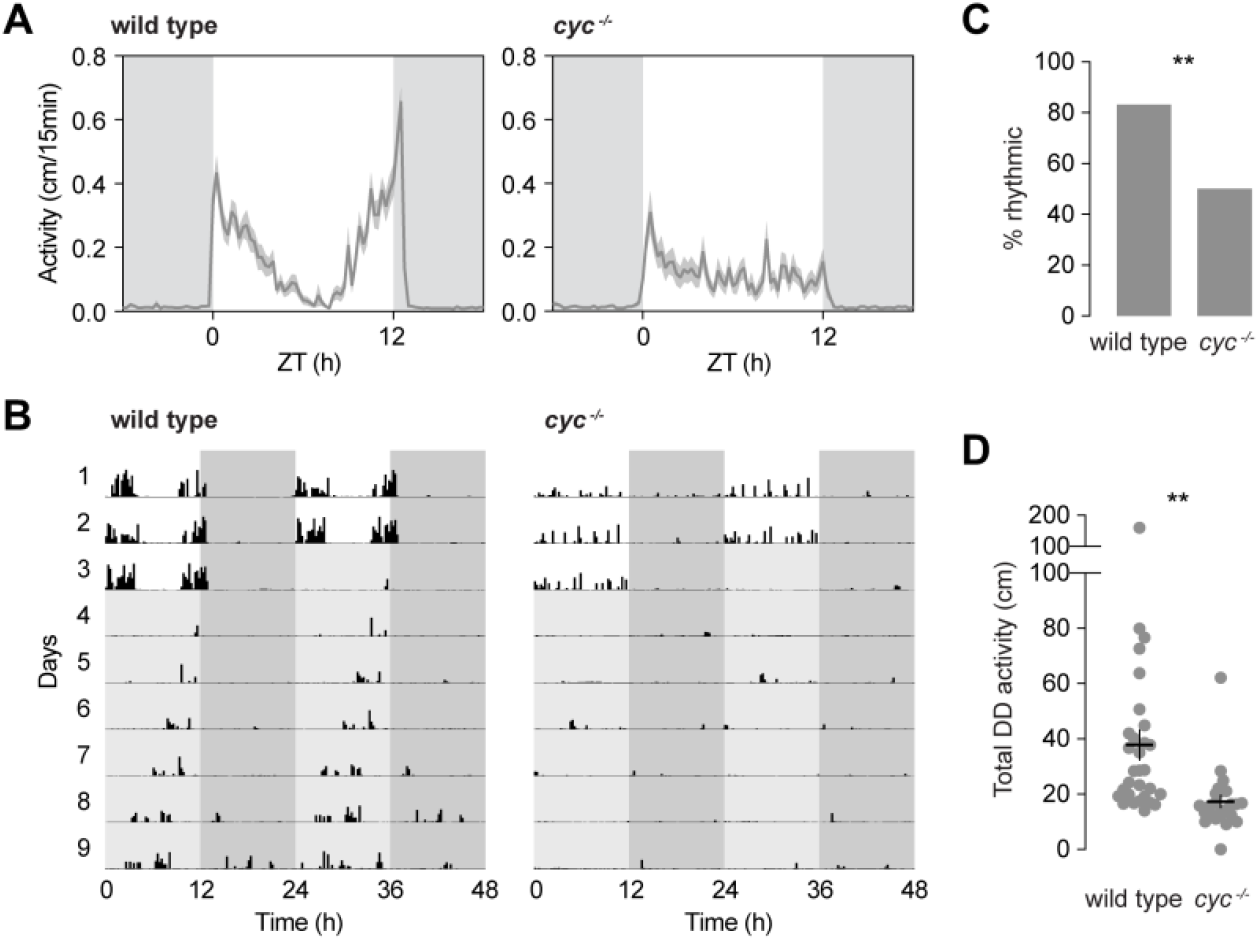
Characterization of locomotor activity in non-blood-fed *cycle* mutants. (A) Daily activity profile of non-blood-fed wild type females (n = 30) and *cycle* mutants (n = 24) under LD entrainment. White and grey shading in represent light and dark phases, respectively. (B) Representative double-plotted actograms of a wild type female and a *cycle* mutant (non-blood-fed). Activity shown in 15-minute bins. Locomotor activity was recorded under three days of 12 hours light: 12 hours dark LD cycles (white and dark grey shading represent light and dark phases, respectively), followed by seven days of DD cycles (light grey shading represents respective light phase from prior LD entrainment). (C) Percentage of rhythmic animals under DD. Animals were considered rhythmic if rhythmic power is greater than 50 (Chi-square periodogram). Chi-square test. (D) Total activity under DD entrainment. Unpaired t-test. Lines and shaded lines in (A) show mean and SEM. Lines and error bars in (D) show mean and SEM and dots show individual animals. **P < 0.01. Exact P values are provided in **Table S1**.

**Figure S3.**
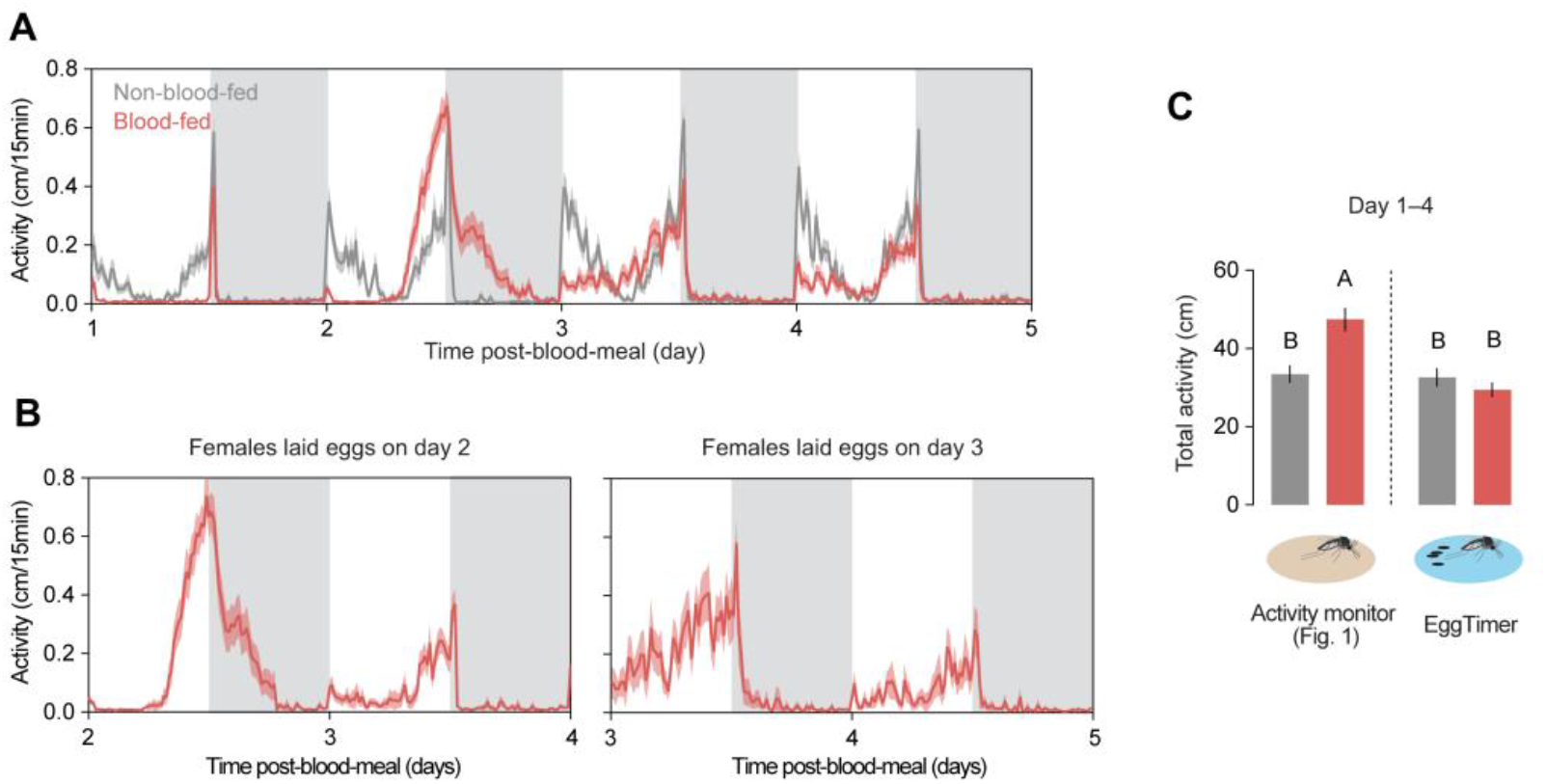
Extended EggTimer recording of wild type females. (A) Locomotor activity profile of non-blood-fed (n = 45) and blood-fed (n = 38) *Aedes aegypti* mosquitoes in EggTimer assay. (B) Locomotor activity profile of blood-fed *Aedes aegypti* mosquitoes in EggTimer assay on egg-laying day and the day after. (C) Total activity days 1–4 post-blood-feeding. Ordinary one-way ANOVA with Tukey’s multiple comparisons test. Data labeled with different letters are significantly different from each other (P < 0.0001). White and grey shading in (A) and (B) represent light and dark phases, respectively. Lines and shaded lines in (A) and (B) show mean and SEM. Bars and error bars in (C) show mean and SEM. Exact P values are provided in **Table S1**.

**Figure S4.**
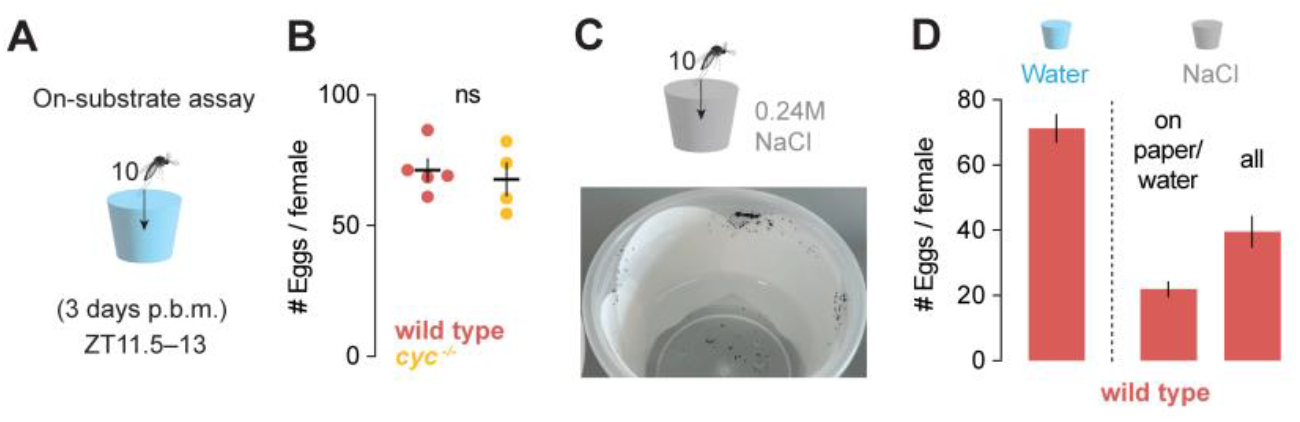
*Cycle* mutants do not show egg-laying deficits when placed directly on egg-laying substrate. (A) Schematic of on-substrate egg-laying assay. Animals three days post-blood-meal (p.b.m.) were loaded in groups of 10 and allowed to lay eggs for 90 mins starting from ZT11.5. (B) Eggs laid per female by wild type (n = 5) and *cycle* mutant (n = 4) *Aedes aegypti* mosquitoes in the on-substrate assay. Unpaired t-test. (C) Representative ovi-cup from saltwater (0.24 M NaCl) on-substrate egg-laying assay. Animals three days post-blood-meal were loaded in groups of 10 and allowed to lay eggs for 90 mins starting from ZT11.5. (D) Eggs laid per female by wild type females in on-substrate assays with water (data from B) and saltwater (n = 6). Since animals avoid saltwater, we report eggs laid on the filter paper and water and all eggs collected (including eggs on the plastic cup). Lines and error bars in (B) show mean and SEM and dots show individual trials. Bars and error bars in (D) show mean and SEM. P > 0.05: not significant (ns). Exact P values are provided in **Table S1**.

